# Investigation of Axonal Beading Induced by Photo-oxidation

**DOI:** 10.1101/2025.02.07.637201

**Authors:** Deepak Mehta, Pramod Pullarkat

## Abstract

Oxidative stress caused by the presence of excessive free radicals (reactive oxygen species) is known for its adverse effects in cellular function. When initiated by fluorescent dye excitation, it can lead to photo-oxidative stress, causing cellular damage. In this study, we investigate axonal beading induced by the photo-oxidation of a membrane based fluorescent dye. Our findings show that apart from the use of oxygen scavengers, chelation of free calcium and stabilizing microtubules or actin filaments by pharmacological agents can mitigate oxidative stress induced axonal beading. Furthermore, we take advantage of this light induced oxidative stress phenomenon to probe axonal response to spatially confined perturbations of varying strength. We demonstrate that low excitation levels lead to long-lived localized beading, whereas higher levels of excitation result in global degeneration and beading of axons. Besides the results presented here, this light based technique is a convenient tool to investigate the effect of local oxidative stress at the tissue level, for example, in cultures of brain slices or organoids.

## 1 Introduction

Axons of the central and peripheral nervous system undergo a wide variety of degenerative processes, resulting in severely debilitating conditions. These include diseases like Alzheimer’s and Parkinson’s, where axonal degradation and atrophy ensues biomolecular triggering events, diabetic or chemotherapy-induced neuropathies, ischemia, edema, and degradation induced by crush or stretch injuries to the nerves. An improved understanding of the degenerative process occurring at the axonal level and the mechanisms that lead to gross axonal destabilization is therefore important for devising strategies to prevent or mitigate damage induced by disease or mechanical insults. Among these mechanisms, the degradation of axons caused by oxidative stress is particularly noteworthy [1–3]. ROS are essential for cell homeostasis and signalling [4] and are produced as metabolic byproducts [5, 6]. However, excessive ROS levels, for example, that result from mitochondrial dysfunction, cause oxidative stress [7–9], and lead to various diseases [10–12] through lipid, protein, or DNA damage. This condition also contributes to the neuronal ageing process [13, 14], and is associated with various neurodegenerative disorders such as Alzheimer’s, Parkinson’s, and Huntington’s. [12, 15].

There are multiple ways of inducing oxidative stress to neurons in culture, like, for example, exposure to nitric oxide [16] [17]. Such induced oxidative stress results in axonal retraction and/or the formation of swellings along the axon, known as axonal beading, which indicate cytoskeletal degradation [14, 18, 19]. Here, we demonstrate that photo-oxidation of an excited fluorescent dye can be used to investigate the pathways of axonal degeneration. We show that the dye in its excited state becomes phototoxic, presumably due to the generation of free radicals that can react with oxygen-forming ROS [20–23]. This induced phototoxicity, in turn, results in the elevation of cytosolic free calcium triggered by the entry of extracellular calcium into the cell, followed by release from internal stores. The increased calcium levels then lead to degradation of the axonal microtubules and actin filaments. Preventing photo-oxidation using a free oxygen scavenger like ascorbic acid, or preventing elevation of cellular free Ca++ using intra/extra-cellular Ca^++^ chelators, or stabilization of either actin filaments or microtubules, all result in suppressing light induced axonal beading. Moreover, we show that localized excitation can be used as a tool to mimic local calcium events generated by injury. The use of light allows us to easily modulate the strength of the local perturbation and allows us to investigate how local damage propagates and affects axonal stability. At a larger scale, as the susceptibility of neurons to oxidative stress shows spatial dependencies *in vivo*, this method may be useful in investigating the effect of local oxidative stress at tissue level using brain slices or brain organoids [24].

## 2 Materials and Methods

### Cell culture

Fertilised Giriraja-2 chicken eggs were obtained from Karnataka Veterinary, Animal and Fisheries Sciences University, Bangalore, India. To culture the chick dorsal root ganglion (DRG) neuronal cells, DRGs were extracted from 9^*th*^ or 10^*th*^ day chick embryo. The DRGs were then treated with Trypsin-EDTA (Gibco, Cat No. 15400-053) for 10 minutes in a calcium- and magnesium-free Hank’s Balanced Salt Solution (HBSS) (Gibco, Cat No. 14175-095). After trypsinization, DRGs were plated in Leibovitz’s (L-15) cell culture media (Gibco, Cat No. 21083-027) containing 0.006 g/ml methylcellulose (ColorconID, Cat No. 34516), mixed overnight at 4 ^°^C, 2% D-glucose (Sigma-Aldrich, Cat No. G6152), 10% Fetal Bovine Serum (Thermo Fisher Scientific, Cat No. 10100–147), 20 ng/ml Nerve Growth Factor (Gibco, Cat No. 13290-010) and 0.5 mg/ml penicillin-streptomycin glutamine (Gibco, Cat No 10378-016). Clean glass coverslips without any additional coatings were used as substrates for cell culture. The cultures were subsequently incubated at 37 ^°^C for a 24-hour period.

### FM4-64 staining

After 24 hours of cell culture, the cells were incubated with FM4-64 (Invitrogen, Cat No T13320) in various concentrations ranging from 2.5 *µ*M to 15 *µ*M as per the need of experiments. A stock solution of FM4-64 at 17 mM was prepared in Dimethyl sulfoxide (DMSO) (Sigma-Aldrich, Cat No D8418), and the desired concentration was added to the cell culture 15 min before the experiment. For the excitation and emission of FM4-64 dye, we used filter sets D535/50x and 565dcxt from Chroma Technology Corp., respectively, at a constant excitation intensity of 0.08 *µ*W/*µ*m^2^, measured using a digital power meter (Thorlab Inc., Part No. PM100D) placed on the microscope stage. The illumination was done using a 100 W HBO lamp.

### Monitoring of intracellular calcium

To measure intracellular calcium, we used Fluo-4, AM at 1 *µ*M (Invitrogen, Cat No. F14201) from a stock of 1 mM prepared in DMSO. A set of Zeiss filters (BP 470/20, FT 493, BP 505-530) was used for the excitation and emission of Fluo-4, AM. The Fluo-4 AM was added to the cell-cultured dish 15 min before the experiment started.

### Calcium chelation

To chelate the extracellular and intracellular calcium, we used EGTA (Ethylene glycol−bis (*β*− aminoethyl ether)−*N, N, N*^*′*^, *N*^*′*^− tetraacetic acid) (Fluka, Cat No 03777-10G) at 5 mM prepared in milli-Q water and BAPTA-AM 1,2-bis(o-Aminophenoxy) ethane -N,N,N’,N’-tetraacetic Acid Tetra (acetoxymethyl) Ester) at 100 *µ*M prepared in DSMO, respectively. Both reagents were added to the cell-cultured dish 15 min before the experiment started.

### Antioxidant treatment

L-Ascorbic acid (Nice, Cat No. A73101) is used for its anti-oxidant behaviour at 4 mM concentration. It was added to the cell culture medium 15 min before fluorescence imaging when photo-oxidation had to be prevented.

### Cytoskeleton stabilizer

To stabilize microtubules and actin filaments, cells are incubated with Taxol (Sigma-Aldrich, Cat No. T7402-1MG) at a concentration of 10 *µ*M and with Jasplakinolide (Invitrogen, Cat No. J7473) at 5 *µ*M for 30 min prior to the experiment.

### Imaging of microtubules

We prepared a stock solution of SPY555-tubulin dye (Spirochrome Inc, Cat No. SC203), by diluting it 1000 times in DMSO. To stain the microtubules, we used 1 *µ*l of the stock solution in 1 ml of cell culture media and incubated the mixture for 1 hour before imaging. Imaging was performed using a Leica TCS SP8 confocal laser scanning microscope with a 561 nm laser.

### Imaging of actin filaments

Neurons were fixed using a solution containing 0.5% (v/v) glutaraldehyde (Sigma-Aldrich, Cat No. G7651-10 ml) and 4% (w/v) paraformaldehyde (Electron Microscopy Sciences, Cat No. 15710) in 1X phosphate-buffered saline (PBS) (Gibco, Cat No. 70011-044) for 10 minutes at room temperature. After fixation, the neurons were rinsed three times with 1X PBS, with a gap of 5 minutes. A solution of 0.45 *µ*M rhodamine phalloidin (Sigma-Aldrich, Cat No. P1951-1MG) along with 0.2% (v/v) Triton X-100 (Sigma-Aldrich, Cat No.T8787-50 ml) was prepared in 1X PBS and added to the fixed neurons for 5 minutes. Following a final wash with 1X PBS, the cells were imaged using a Leica TCS SP8 confocal laser scanning microscope with a 561 nm laser.

### Microscopes used for imaging

For imaging, we employed a Zeiss AxioObserver Z1 inverted microscope equipped with a 40x/0.5 Ph2 phase contrast objective. Data was collected using an Axio Cam MRm CCD camera through AxioVision SE64 software, and analysis was conducted using ImageJ. Additionally, we utilized a Leica TCS SP8 confocal laser scanning microscope for the localized excitation of the FM4-64 dye, using a 488 nm laser at a power of 6 mW. The Fluorescence Recovery After Photobleaching (FRAP) tool was applied with the 40x/0.5 Ph2 phase contrast objective. FRAP allows for excitation over a 5 *µ*m segment in the middle of the axon. All experiments were conducted at 37 ^°^C.

## 3 Results and Discussion

### 3.1 Photo-oxidation of FM4-64 dye causes axonal beading

The experimental procedure is detailed schematically in Figure 1A. One day *in vitro* (DIV1) neurons from chick dorsal root ganglia were incubated with FM4-64 fluorescent dye at a concentrations of 2.5 *µ*M, 5 *µ*M, 10 *µ*M or 15 *µ*M, for 15 minutes at 37 ^°^C. At time t = 0 minutes, the dye was excited for 1 minute, maintaining a constant excitation intensity of 0.08 *µ*W/*µ*m^2^. Fluorescence images were acquired during this period. At time t = 1 minute, the excitation of FM4-64 was stopped, and phase contrast imaging was started to monitor axonal beading. Figure 1B shows a time sequence of phase contrast and fluorescence images of an axon labeled with 10 *µ*M dye (also see Suppl. Movie S1). It can be seen that the initially tubular axon develops a series of nearly axisymmetric swellings its entire length post-exposure. This shape change, often referred to in the literature as axonal beading, progresses after the excitation light is turned off. This axonal beading was non-reversible during up to the maximum observation time of 30 min.

**Figure 1:**
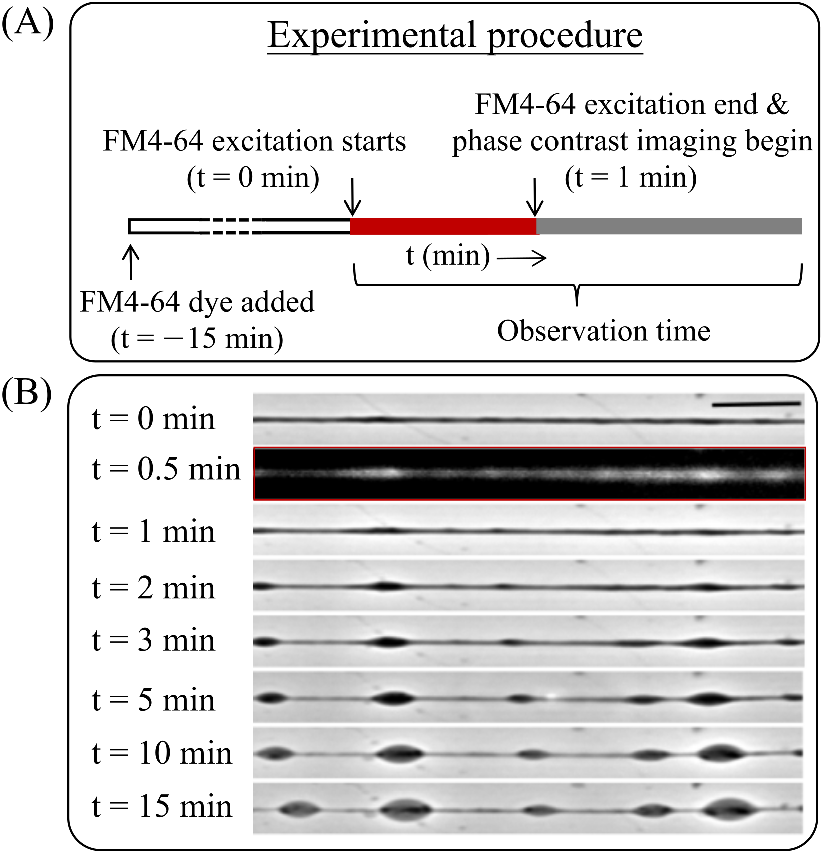
Experimental procedure and light induced axonal beading. **(A)** Chick dorsal root ganglia axons were pre-incubated with FM4-64 fluorescent dye for 15 minutes. Then, at time t = 0 min, the dye was excited for 1 minute. Fluorescence images were acquired during this period. At t = 1 min, the excitation light was turned off, and imaging was switched to phase contrast mode to observe the evolution of axonal shape. **(B)** Sequence of images showing the time evolution of an axon which was illuminated by the FM4-64 excitation light for one minute. The second image in the sequence (t = 0.5 min) shows the fluorescence from the dye during illumination, and the rest are phase contrast images. Beading progresses even after the excitation light is turned off. Note the highly axisymmetric nature of the axonal beading (also see Suppl. Movie S1). The scale bar represents a length of 10 *µ*m.

When the concentration of the FM4-64 dye is increased in steps from 2.5 *µ*M to 15 *µ*M, we observe that the beading time–the duration taken for bead-like structures to appear along the axon–decreases as shown in Figure 2A. Specifically, the beading times (mean *±* standard error of the mean) for FM4-64 concentrations of 2.5, 5.0, 10.0, and 15.0 *µ*M, were 4.16 *±* 0.40 min, 2.86 *±* 0.16, 2.55 *±* 0.15 min, and 1.83 ± 0.12 min, respectively. To determine whether the observed beading was due to DMSO, which was used as a solvent for the dye, we tested the cells with up to 5%v/v DMSO and without the FM dye on 19 axons, and none exhibited beading upon exposure to the same intensity and duration of the excitation light. Additionally, we used three different batches of FM4-64 dye (Lot No. 33034W, Lot No. 2140304, and Lot No. 11452558) to check for consistency in beading. All three batches induced beading in the axons with similar timescales. We then examined 25 axons labelled with FM4-64 at a concentration of 5 *µ*M and watched the neuronal cells without using excitation light to determine if beading occurred due to the dye’s intrinsic toxicity. None of the observed axons exhibited beading in the absence of excitation of the dye. Based on these observations, we went on to investigate the possibility of photo-oxidation caused by dye excitation.

**Figure 2:**
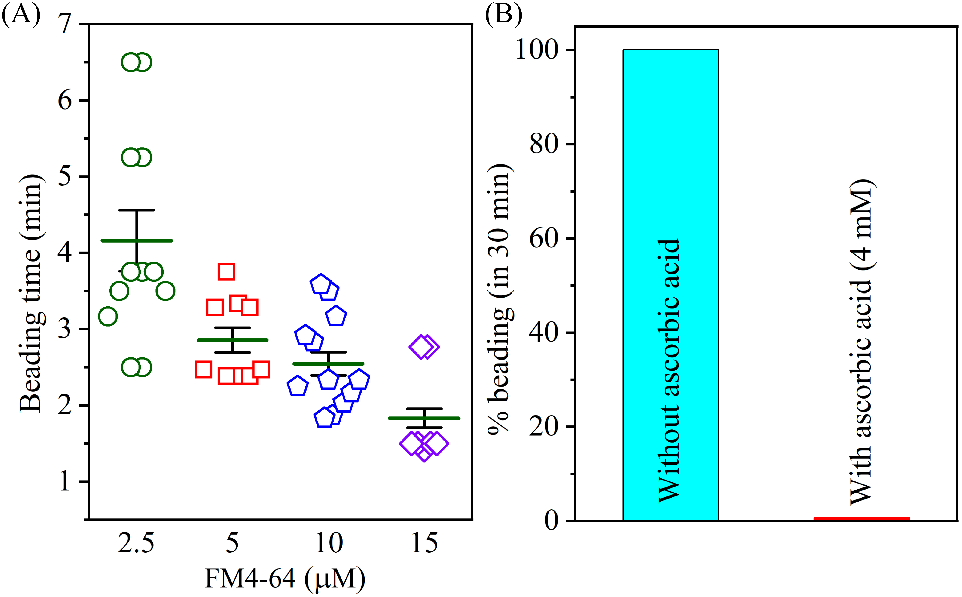
Changes in beading time due to FM4-64 concentration or antioxidant treatment. (**A**) The time taken for beads to appear decreases monotonically with increasing dye concentration. Axons were illuminated at a constant excitation intensity of 0.08 *µ*W/*µ*m^2^ for a period of 1 min. The measurements are for 2.5 *µ*M, 5 *µ*M, 10 *µ*M and 15 *µ*M of FM4-64, with each data point corresponding to one axon. The corresponding average beading time is represented using a green horizontal line, and the error bars are standard mean errors. We note that while performing the measurement, a few axons did not show beading even over the experimental timescale of 15 min. We observed that 12/16, 16/16, 14/15, 14/17 and 16/18 axons showed beading for FM4-64 concentrations of 2.5, 5.0, 10.0, and 15.0 *µ*M, respectively. The error bars are the standard error of the mean. **(B)** Bar plots showing the effect of 4 mM ascorbic acid on beading for axons labeled with 5 *µ*m FM4-64. The intensity and duration of excitation used were the same as for the previous cases. All 16 axons that were observed exhibited beading when no ascorbic acid was added. However, in ascorbic acid containing medium, out of 25 axons observed, none of the axons showed beading.

To investigate the potential for photo-oxidation arising from FM4-64 excitation, we pretreated the cells with 4 mM L-ascorbic acid as an antioxidant. In this case, axons labelled with 5 *µ*M FM4-64 and subjected to the same excitation protocol did not exhibit any beading up to 30 min of observation time, and the data is shown in Figure 2B. From this, we can conclude that photo-oxidation effects are responsible for axonal beading.

### 3.2 Photo-oxidation drives Ca^++^ influx leading to axonal beading

Next, we investigated changes to intracellular free calcium induced by photo-oxidation using axons labeled with FM4-64 (5 *µ*M) and Fluo4 AM (1 *µ*M). For imaging, the FM4-64 membrane dye was excited for 1 min as before, and the free Ca^++^ levels were monitored just before FM4-64 excitation and then every 0.5 min by switching the microscope filter sets. These observations revealed a 5-6 fold increase in intracellular calcium levels following excitation of FM4-64. The inset to Fig. 3A illustrates the variations in free Ca^++^ in an axon at distinct time points. A sharp increase in Ca^++^ can be seen post-excitation. The localized higher intensity of calcium fluorescence at the beads is due to the higher cytoplasmic volumes of the beads as compared to the thinned-out regions connecting adjacent beads (see Suppl. Figure S1). As can be seen from Fig. 3A, a significant increase in Ca^++^ levels began at about 0.5 min of FM4-64 excitation and continued to increase even after the membrane dye excitation was stopped at 1 min.

**Figure 3:**
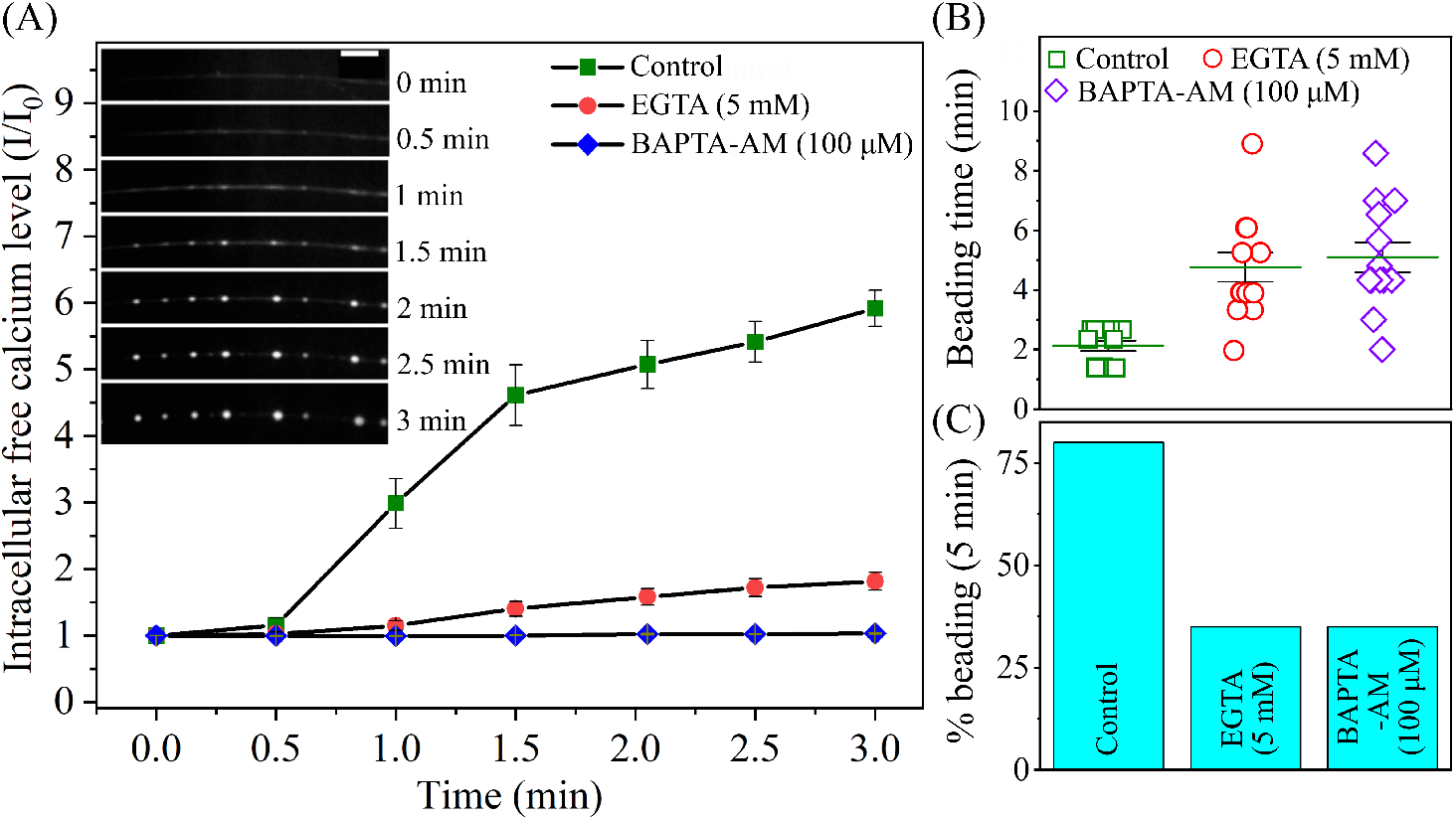
Calcium chelation suppresses photo-oxidation-induced axonal beading. **(A)** The inset shows the time evolution of the fluorescence from the intracellular Ca^++^ indicator Fluo4-AM after the FM4-64 dye is excited (t = 0 min). The scale bar is 10 *µ*m. To quantify changes in axonal Ca^++^, the intensity (*I*) integrated over the length of the entire axonal segment was normalized with its pre-excitation intensity (*I*_0_). The green squares show the change in intracellular calcium for the control axons, which were labeled with FM4-64 and Fluo4 AM (n = 10). The red circles and the blue diamonds show the changes in intracellular calcium for the double-labeled axons pre-treated with 5 mM EGTA (n = 9 axons) and 100 *µ*M BAPTA-AM (n = 7 axons), respectively. The scale bar indicates a length of 10 *µ*m. **(B)** The plot shows the beading time of axons after FM4-64 excitation for control axons (n = 13) and for axons that were pre-treated with EGTA (5 mM) (n = 13), and those treated with BAPTA-AM (100 *µ*M) (n = 12). **(C)** The bar plot shows the percentage of beaded axons after FM4-64 excitation for the control axon (n = 20) and when the axons were pre-treated with EGTA (n = 20) and BAPTA-AM (n = 20) as in previous examples. Error bars are standard errors of the mean.

In order to see if axonal beading was caused by elevated Ca^++^ levels, we chelated extra- and intracellular calcium using 5 mM EGTA and 100 *µ*M BAPTA-AM, respectively. As can be seen from Fig. 3A, these treatments drastically decreased the free calcium change seen after FM4-64 excitation. A correlated increase in beading time and a corresponding decrease in the number of beaded axons was also observed in calcium-chelated cells, suggesting that Ca^++^ suppression leads to suppression of beading (see Fig. 3B, C).

### 3.3 Stabilization of microtubules or actin filaments mitigates photo-oxidation damage

To investigate the role of the axonal cytoskeleton in photo-oxidation-induced axonal beading, we treated cells with either 10 *µ*M of Taxol, which stabilizes microtubules or 5 *µ*M of Jasplakinolide, which stabilizes actin filaments. Such treatments resulted in a significant reduction in the percentage of beaded axons post-photo-oxidation of the membrane dye. As can be seen from Fig. 4, 25 out of 33 control axons (FM4-64 labeled) exhibited beading within 5 minutes. In contrast, only 5 out of 49 axons showed beading when pre-treated with Taxol, and only 8 out of 50 axons exhibited beading following pre-treatment with Jasplakinolide. When axons were co-treated with both Taxol and Jasplakinolide, only 2 out of 44 axons displayed beading. These results suggest that phototoxicity-induced Ca^++^ increase leads to cytoskeletal damage, which in turn causes axonal beading.

**Figure 4:**
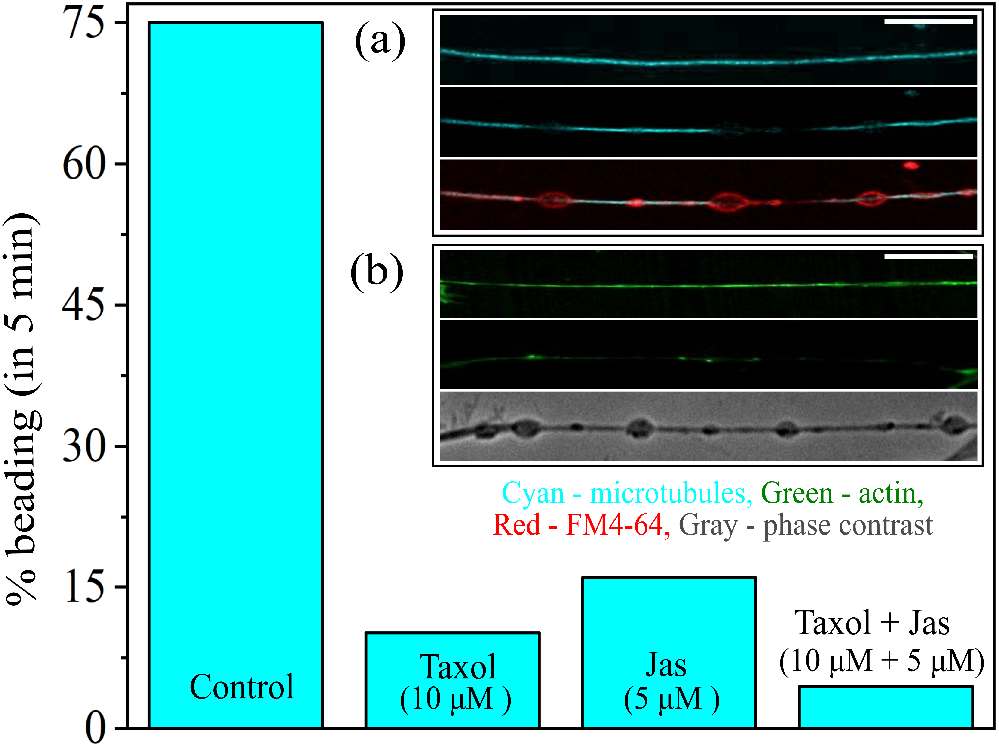
Cytoskeleton stabilization suppresses axonal beading. The bar plot shows the percentage of beaded axons within 5 minutes of photo-oxidation for control axons (n = 33), axon pretreated with 10 *µ*M of Taxol (n = 49), axons pre-treated with 5 *µ*M of Jasplakinolide (n = 50), and axon pre-treated with Taxol (10 *µ*M) and Jasplakinolide (5 *µ*M) together (n = 44). **(Inset)** Confocal images of axons labeled with either the microtubule dye SPY555-tubulin or with rhodamine-phalloidin that labels actin filaments. **(a)** From top to bottom: fluorescence images showing microtubules labelled with SPY555-tubulin in an axon before the FM4-64 dye was excited, the image of the same axon obtained in the SPY555 channel after the axon has beaded due to excitation of the membrane dye, and the FM4-64 channel showing the axon shape. **(b)** Top image shows fluorescence from actin filaments in an axon, which was fixed and labelled with rhodamine-phalloidin before excitation. The next image shows a different axon that was labelled with rhodamine after excitation and formation of beads. The FM4-64 channel does not reveal membrane shape in this case because the cells were permeabilized for labelling. Hence, a phase contrast image of the same beaded axon taken before permeabilization is shown as the bottom image (see Suppl. Figure S2 for more examples). The scale bar indicates 10 *µ*m.

To test this further, we imaged microtubules using SPY555-tubulin dye and actin filaments using rhodamine-phalloidin in control cells. As can be seen from the inset of Fig. 4, widespread depolymerization of microtubules and actin is observed along the axonal segment post-excited.

### 3.4 Changes to axonal volume post-excitation

The light induced photo-oxidation, the release of free calcium, and the subsequent degradation of the cytoskeleton and other components could change the intra-cellular osmolarity and water permeability across the membrane. To check for such effects, we measured the evolution of axonal volume using a home-developed Matlab code. This method takes advantage of the axisymmetric shapes of the axon before and after beading and computes the volume from the two automatically detected axonal interfaces (see Suppl. Figure S3). As the local diameter varies along the axon, the volume is computed for segment lengths that cover an integral number of beads. The evolution of volume, normalized by the initial volume, is shown in Fig. 5. This data clearly shows that axonal volume increases with time post-excitation.

**Figure 5:**
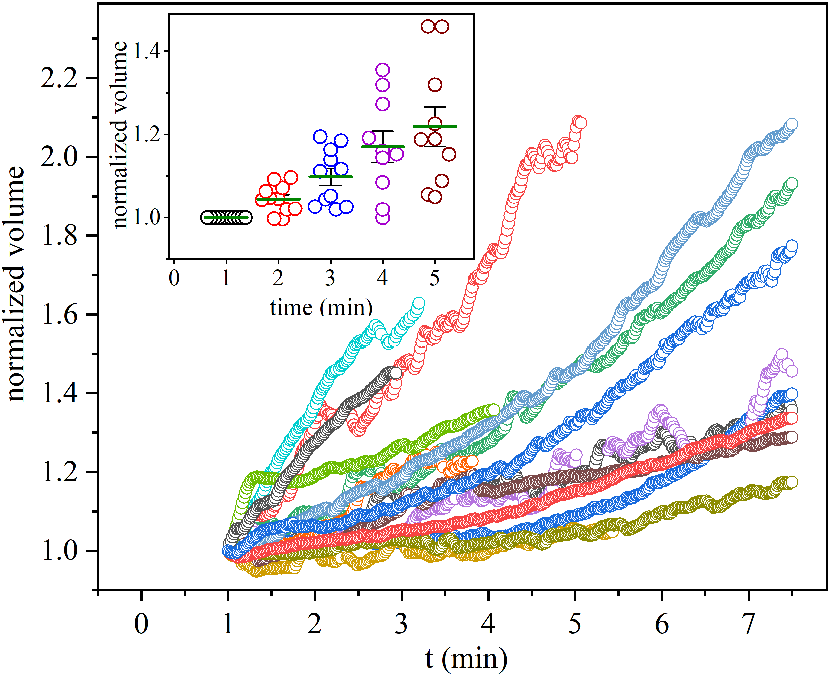
Volume evolution post-excitation. Axonal volume was determined using image processing and considering the shapes to be axisymmetric at all times. The main plot shows the time evolution of the normalized volume of individual axons (n = 15) from the time excitation stopped. The normalized volume was computed as V(t)/V(t = 1 min). The inset shows the the values for a few time points, and the averages indicate a linear increase in volume within the observation time.

### 3.5 Axonal beading in response to local photo-oxidation

The use of light to induce photo-oxidative damage to axons allows us to use it as a tool to investigate the resilience of axons to local damage and the resulting Ca^++^ elevation. For this, we used a confocal microscope’s Fluorescence Recovery After Photobleaching (FRAP) tool to excite FM4-64 dye locally over a 5 *µ*m segment of the axon. Local photo-oxidation of FM4-64 dye (5 *µ*M) resulted in a spatiotemporal increase in intracellular calcium levels. Confocal imaging conducted on cells co-labelled with Fluo4-AM revealed a gradual rise in free Ca^++^ originating from the excitation site and propagating bidirectionally along the axon as pictured in Fig. 6A. A quantitative analysis supported this observation, demonstrating a progressive increase in normalized Fluo4-AM fluorescence intensity as a function of distance from the excitation centre at different time points following photo-oxidation and is illustrated in Fig. 6B. We observed this spread in free calcium level in all 9 axons that were studied. Morphological changes accompanied this Ca^++^ wave with a lag time, as can be seen from Fig. 6A (also see Suppl. Movie S2). An estimate of the speed of the calcium wave can be made from the images by applying a threshold to the fluorescence intensity to define a propagating front. It is seen that the axonal Ca^++^ wave propagates at a speed of 39.6 *±* 6.3 *µ*m/min (9 axons)(data in (**B**) Bar plots displaying the number of axons with multiple beads (green bars), no beads (red bars), and a single bead (purple bars) for 1.5 min and 3.0 min excitations of FM4-64.

**Figure 6:**
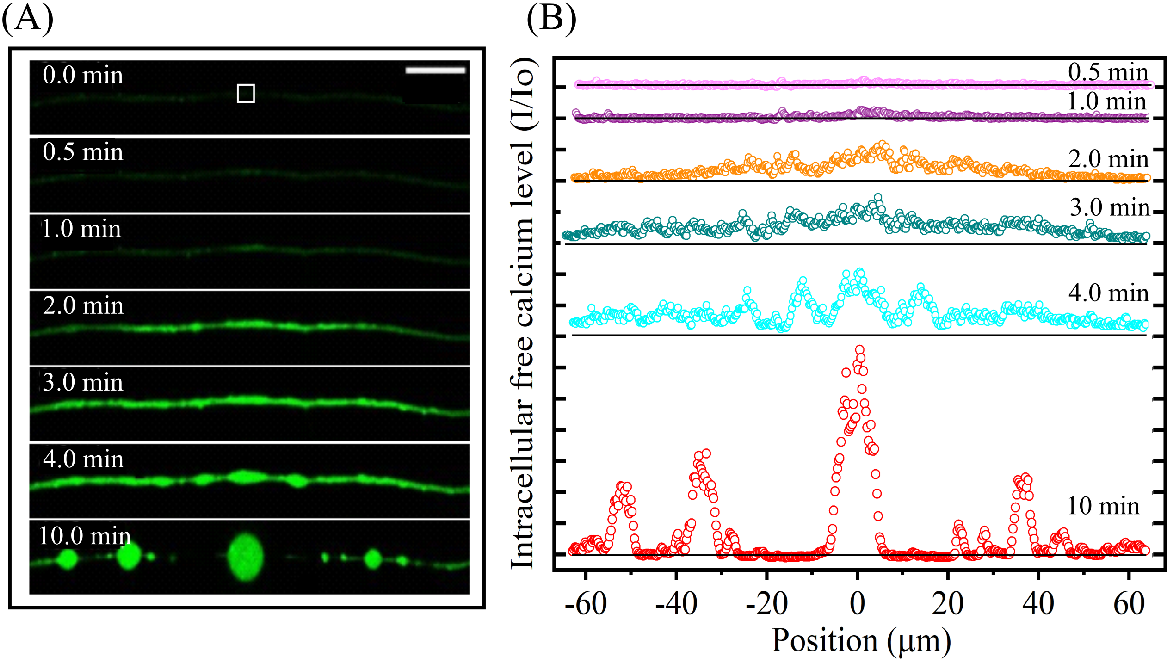
Spatiotemporal evolution of beading after local photo-oxidation. **(A)** A sequence of confocal images showing the intracellular calcium levels and beading in an axon subjected to local photo-oxidation. The sale bar is 10 *µ*m. At time t = 0 min, the axon, labelled with FM4-64 (5 *µ*M), is excited over a 5 *µ*m segment in the middle (white square). An increase in intracellular calcium levels is observed over time, as illustrated by snapshots taken at t = 0.5 min, 1.0 min, 2.0 min, 3.0 min, 4.0 min, and 10.0 min. The calcium levels increased from the excitation region and gradually spread bidirectionally in every case (n = 9). **(B)** Plots of the Fluo4-AM fluorescence intensity, normalized to its pre-excitation value, as a function of position along the axon from the excitation point (same axon shown in (A)). The black horizontal lines represent the pre-excitation intensity providing a reference to compare the intensity at subsequent time points with the initial value.

Suppl. Figure S4). To compare this speed with another form of axonal perturbation, we subjected axons to mechanical stretch using a glass needle. In such cases (n = 10), the entire axon experiences mechanical stretch and calcium spreads across the entire axonal segment that was imaged (≃ 85 *µ*m) in less than a second (measurement of speed limited by the acquisition speed of 1 fps), (see Suppl. Movie S3).

Next, we explored how the strength of the local light induced perturbation affects axonal stability. An excitation of 1.5 min of FM4-64 at a concentration 5 *µ*M primarily resulted in the formation of a single bead in the vicinity of the excitation site, as illustrated in Fig. 7A(i). In contrast, prolonged photo-oxidation of 3.0 min, with the same dye concentration, led to the appearance of multiple beads distributed along the axon, as shown in Fig. 7A(ii). We quantified these observations by counting the number of axons exhibiting a single bead, those showing multiple beads, and those showing no beading at all following local excitation (see Fig. 7B).

**Figure 7:**
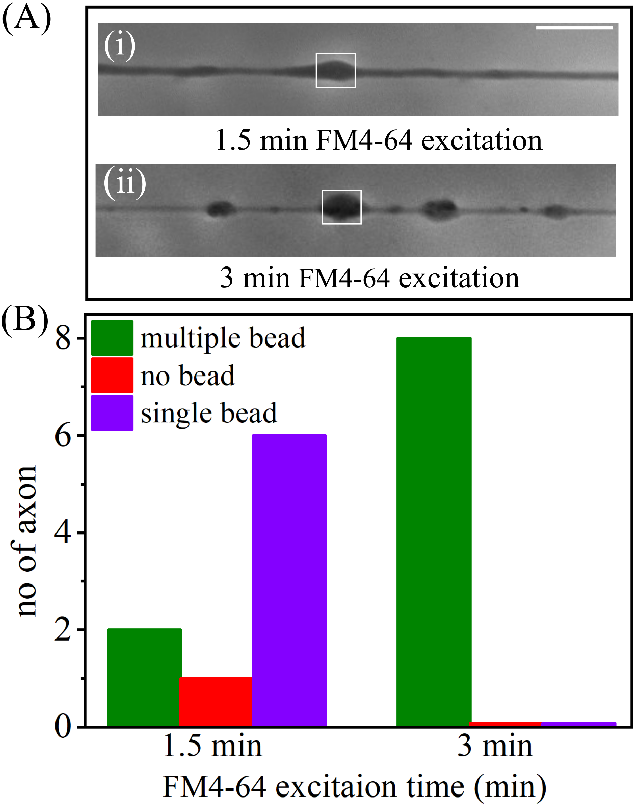
Axonal beading in response to local photo-oxidation. **(A)** Phase contrast images of an axon that was excited only across a 5 *µ*m segment (white box) for a duration of 1.5 min (image i) and one that was excited for 3.0 min (image ii). Only a single bead formed in the case of the axon with short excitation and there was no recovery up to the observation time of 15 min. For longer excitations within the same 5 *µ*m window, multiple beads evolved along the axon away from the excited region (also see Suppl. Movies S4 and S5). The scale bar represents a length of 10 *µ*m.

## 4 Discussion

In this study, we have systematically examined the role and consequence of photo-oxidation caused by the excitation of the fluorescent dye FM4-64. We observe that axons undergo light induced beading–a shape transformation usually seen in degenerating axons. The sequence of events that leads to beading is as follows. Excitation of the dye causes oxidative stress resulting from photo-oxidation. This is evident from the fact that treatment of cells with ascorbic acid suppressed axonal beading. This initial perturbation leads to a sharp elevation of cytosolic free Ca^++^. Treatment with EGTA, a chelator of extra-cellular Ca^++^, suppresses light induced beading, suggesting that entry of extra-cellular calcium into the cell is essential. Treatment with BAPTA-AM, a cell-permeable Ca^++^ chelator, which does not affect extra-cellular calcium, too suppresses Ca^++^ elevation and beading. This suggests that Ca^++^ release from internal stores forms a significant part of the excess free Ca^++^ generated by the action of light. The elevated Ca^++^ levels lead to the degradation of the axonal cytoskeleton. This is evident from the fact that stabilizers of actin filaments or microtubules suppresses light induced beading. Moreover, fluorescence imaging of actin filaments and microtubules show that these filaments are disrupted in regions exposed to light. Phototoxicity also induces an increase in axonal volume, presumably due to increased internal osmolarity.

The beading transformation reported here provides information on the possible mechanisms for the shape change. It has been proposed that beading can occur as a result of interrupted axonal traffic resulting from induced stress or stretch-induced microtubule damage [14, 25–30]. As can be seen from the images presented here, the observed shapes we see are highly axis-symmetric. Such shape transformations have been studied extensively in synthetic tubular lipid bilayer membranes that are subjected to mechanical stress [31, 32] and in other soft elastic cylindrical materials [33]. It has been shown that axons, too, can undergo such membrane/interfacial tension-driven shape transformations under hypo-osmotic conditions [34, 35]. The increase in volume we observe in our case suggests that a similar mechanism could be at play. Axonal shape transformations can also be induced by various cytoskeletal perturbations [18, 36, 37]. It has been argued that the stability of the cylindrical shape of an axon depends on the balance between the tension on the axonal membrane and the elastic integrity of the internal cytoskeletal ultrastructure [18]. We observe that the light induced beading is suppressed in the case of axons exposed to stabilizers of either microtubule or actin filament. This suggests that cytoskeletal degradation is the main driving mechanism for phototoxicity or oxidative stress induced axonal beading reported here. The increase in axonal volume is likely a secondary effect, driven by increased intra-cellular osmolarity due to cytoskeletal disruption leading to the generation of monomers and the release of sequestered molecules.

The use of light as a tool to induce oxidative stress allows us to explore the response of axons when oxidative stress is applied locally, which is not possible using standard methods. At low excitation time, only a single bead forms at the perturbed location when only a 5 *µ*m segment is excited. For longer excitation times, beads form progressively away from the excited region. A wave of free calcium propagates at a speed of about 40 *µ*m/min. Beading lags behind calcium elevation, and extensive cytoskeletal damage is also observed. These observations suggest that cytoskeletal damage results from elevated calcium levels. Such experiments provide information on the modes of axonal damage when subjected to local injury causing conditions that result in Ca^++^ based degradation events, for example, crush injuries to nerves [28, 29, 38]. Interestingly, it has been suggested recently that axonal beading may have a protective role by hindering calcium wave propagation [39]. Moreover, the experiments presented here suggest that actin filament and microtubule stabilizers, especially Paclitaxel, which is currently in use as an anti-cancer drug [40], are potential pharmacological intervention strategies to mitigate Ca^++^ driven axonal damage, for example, crush injuries to nerves. Recently, microtubule tip binding proteins have been suggested as a potential target for stabilizing axons against beading transformation [14].

The ability of our optical method to apply oxidative stress to individual neurons, or locally along axons, and the ease of adjusting the strength of oxidative perturbation by adjusting the intensity or duration of exposure make this method a versatile tool to study axonal degeneration under oxidative perturbations. It will be interesting to extend this technique to investigate degenerative processes in cultures of brain slices or brain organoids, where susceptibility to damage varies spatially, and local stresses may propagate within the brain tissue [41]. Understanding the axonal damage mechanisms and local vs global effects is critical in applications like photodynamic therapy, which is one of the strategies to treat brain tumors [42]. Such experiments, along with investigation of the mechanisms of bead formation, are also highly relevant in understanding conditions like ischemia [43–45]. In addition, the ability to induce local Ca^++^ events in a controlled manner provides us with an easy technique to study Ca^++^ wave propagation in axons and to investigate the degeneration mechanisms of local injury to nerves [29, 38]. Thus, apart from the mechanisms presented here, the technique we have developed could find multiple applications for the investigation of neurodegeneration.

## Acknowledgements

The authors acknowledge support through The Wellcome Trust DBT India Alliance (grant IA/TSG/20/1/600137)

