## Supplementary information for "Investigation of Axonal Beading Induced by Photo-oxidation"

#### 1 FM4-64 dye

[Fei-Mao (N− (3−triethylammoniumpropyl)−4− (6− ( 4− (diethylamino)phenyl) hexatrienyl ) pyridinium dibromide)]. This is a lipophilic styryl dye belonging to the FM dye family [1], widely used for fluorescence imaging of plasma membranes and to track vesicles in neurons [2, 3]. FM4-64 molecules consist of a lipophilic tail and lipophobic head. The lipophilic tail inserts it into the membrane's outer leaflet, whereas the lipophobic head prevents it from crossing the membrane. Its high quantum yield when it is incorporated a membrane compared to the free dye molecules in the cytoplasm or culture medium makes it an excellent membrane marker. The risk of dye toxicity to cells increases with its improper storage, improper stock concentrations, higher concentrations in samples, and increased light exposure while imaging. [4, 5].

#### 2 Supplementary figures

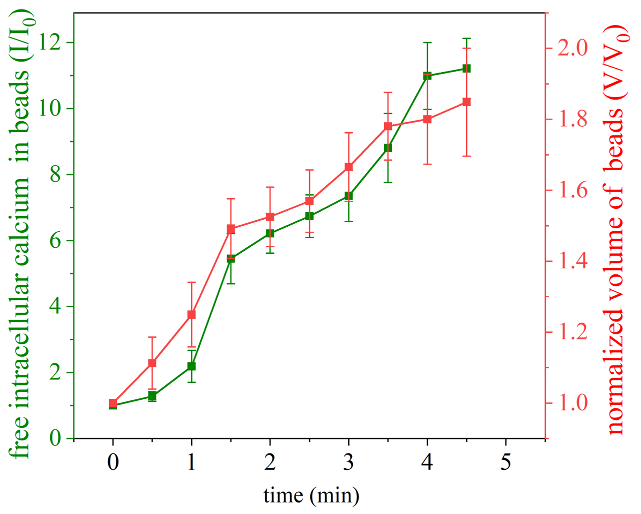


Figure S1: The plot illustrates the ratio of free intracellular calcium levels, denoted as I(t)/I(t = 0 min), as well as the change in normalized volume, V(t)/V(t = 0 min), of beads (n = 10 beads) during photo-oxidation. The correlation shows that the beads have a higher localized intensity of calcium fluorescence due to their larger cytoplasmic volume compared to the thinner regions connecting the adjacent beads.

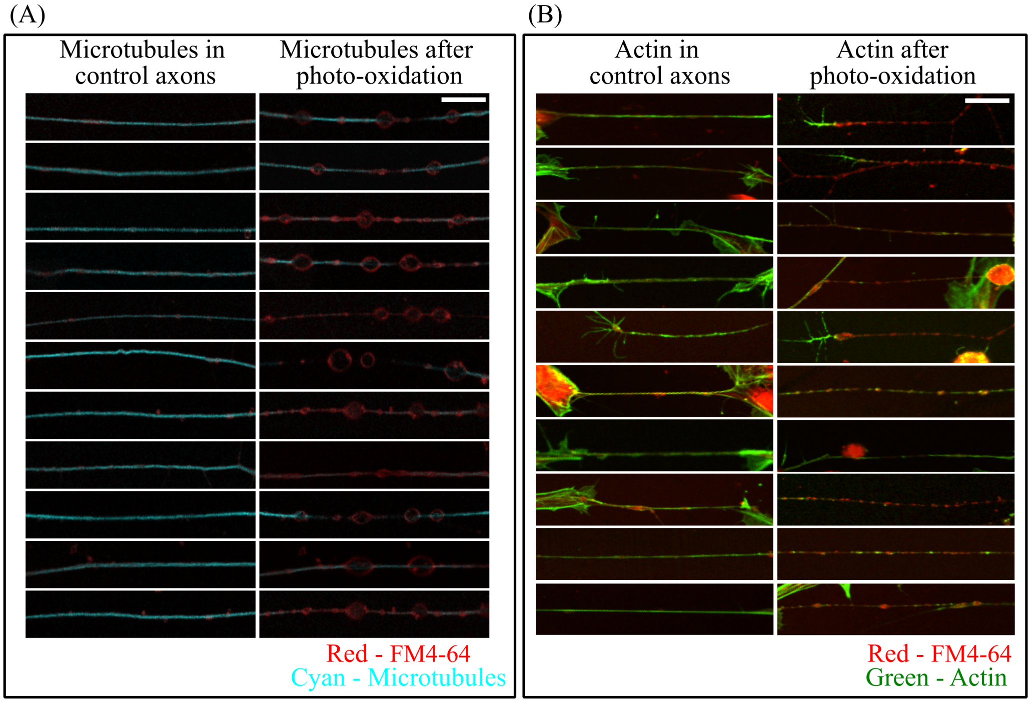


Figure S2: **Cytoskeleton damage post-photo-oxidation.** **(A)** The left images illustrate the combined fluorescence of microtubules labelled with the live cell marker labelled with SPY555-tubulin (cyan) and FM4-64 membrane dye (red) in multiple control axons. The images to the right of panel A show the same set of axons in the same sequence after photo-oxidation. Note the widespread disruption of microtubules in these axons. The scale bar indicates 10 $\mu$m. **(B)** The images on the left show the combined fluorescence images of actin filaments labelled with rhodamine-phalloidin dye (green) and FM4-64 membrane dye (red) in multiple unperturbed axons. The axons were fixed and permeabilized for actin filament labelling. The images on the right of panel B show different photo-oxidized axons. Much less actin filaments are visible in these cases compared to non-excited axons. The scale bar indicates 10 $\mu$m.

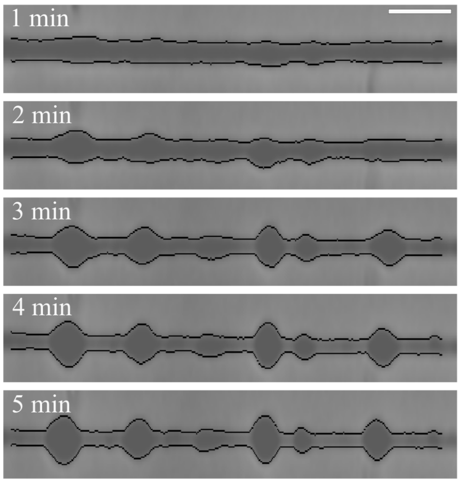


Figure S3: Images showing the analysis of volume during axonal beading. Axonal boundaries are identified based on the intensity gradient across the interface in the radial direction. Points of maximum gradient are traced along each edge of the axon (shown in black). Volume is then computed by assuming axi-symmetry. Images correspond to time points of t = 1 min, 2 min, 3 min, 4 min, and 5 min. The analysis code was written using Matlab.

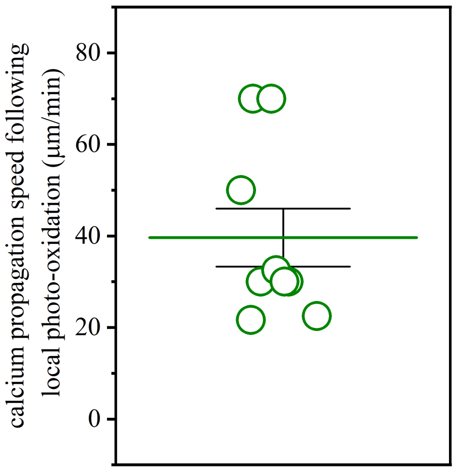


Figure S4: The plot shows the speed of calcium wave induced by local photo-toxicity (n = 9 axons). The green horizontal line is the average and the error bars are standard error of the mean.

### 3 Supplementary movies

1. **Movie S1. - Timelapse showing axonal beading after photo-oxidation.**

   **File name:** photooxidation_induced_axoanl_beading.mp4

   **Movie details:** At time t = 0 min, an axon labelled with FM4-64 dye was excited for 1 minute. At t = 1 min, the excitation light was turned off, and imaging was switched to phase contrast mode to observe the evolution of axonal shape. Note the significant photobleaching seen during the excitation window.

2. **Movie S2. - The slow spread of intracellular calcium during local photo-oxidation.**

   **File name:** slow_spread_of_calcium_during_photooxidation.mp4

   **Movie details:** The bidirectional propagation of cytoplasmic free calcium along the axon seen after local photo-oxidation over a 5 $\mu$m segment (white box in video) at the axon's midpoint.

3. **Movie S3. - The fast spread of intracellular calcium during mechanical damage using a micro glass needle.**

   **File name:** fast_spread_of_calcium_during_mechanical_stretch.mp4

   **Movie details:** An axon is subjected to sudden mechanical stretch using a glass needle. The entire axon experiences mechanical stretch, and calcium spreads across the entire axonal segment in less than a second.

4. **Movie S4. - Single bead formation for low excitation time.**

   **File name:** single_bead_formation_with_low_excitation_time.mp4

   **Movie details:** An axon labelled with FM4-64 was locally excited (white box in video) for a duration of 1.5 minutes. Phase contrast imaging began at t = 1.5 minutes to observe the evolution of the axon's shape. This low excitation time leads to the formation of a single bead near the excitation site.

5. **Movie S5. - Multiple beads for higher excitation time.**

   **File name:** multiple_bead_formation_with_high_excitation_time.mp4

**Movie details:** An axon labelled with FM4-64 was locally excited (white box in video) for a duration of 3 minutes. Phase contrast imaging began at t = 3 minutes to observe the evolution of the axon's shape. This prolonged excitation time leads to the formation of multiple beads along the axon.
